# Automated Isoform Diversity Detector (AIDD): A pipeline for investigating transcriptome diversity of RNA-seq data

**DOI:** 10.1101/2020.01.22.915348

**Authors:** Noel-Marie Plonski, Emily Johnson, Madeline Frederick, Heather Mercer, Gail Fraizer, Richard Meindl, Gemma Casadesus, Helen Piontkivska

## Abstract

**Background:** As the number of RNA-seq datasets that become available to explore transcriptome diversity increases, so does the need for easy-to-use comprehensive computational workflows. Many available tools facilitate analyses of one of the two major mechanisms of transcriptome diversity, namely, differential expression of isoforms due to alternative splicing, while the second major mechanism - RNA editing due to post-transcriptional changes of individual nucleotides – remains under-appreciated. Both these mechanisms play an essential role in physiological and diseases processes, including cancer and neurological disorders. However, elucidation of RNA editing events at transcriptome-wide level requires increasingly complex computational tools, in turn resulting in a steep entrance barrier for labs who are interested in high-throughput variant calling applications on a large scale but lack the manpower and/or computational expertise.

**Results:** Here we present an easy-to-use, fully automated, computational pipeline (Automated Isoform Diversity Detector, AIDD) that contains open source tools for various tasks needed to map transcriptome diversity, including RNA editing events. To facilitate reproducibility and avoid system dependencies, the pipeline is contained within a pre-configured VirtualBox environment. The analytical tasks and format conversions are accomplished via a set of automated scripts that enable the user to go from a set of raw data, such as fastq files, to publication-ready results and figures in one step. A publicly available dataset of Zika virus-infected neural progenitor cells is used to illustrate AIDD’s capabilities.

**Conclusions:** AIDD pipeline offers a user-friendly interface for comprehensive and reproducible RNA-seq analyses. Among unique features of AIDD are its ability to infer RNA editing patterns, including ADAR editing, and inclusion of Guttman scale patterns for time series analysis of such editing landscapes. AIDD-based results show importance of diversity of ADAR isoforms, key RNA editing enzymes linked with the innate immune system and viral infections. These findings offer insights into the potential role of ADAR editing dysregulation in the disease mechanisms, including those of congenital Zika syndrome. Because of its automated all-inclusive features, AIDD pipeline enables even a novice user to easily explore common mechanisms of transcriptome diversity, including RNA editing landscapes.

## Background

Transcriptome complexity and diversity, including patterns of differential isoform expression, non-canonical transcripts, diversity of non-coding RNAs, and regulation of RNA editing play fundamental roles in both normal physiological function and disease mechanisms (ENCODE_Project_Consortium 2004; Albert and Kruglyak 2015; Ardlie and Guigo 2017; Gallo et al. 2017). Due to advances in deep sequencing technologies, RNA-seq experiments have become a more affordable and therefore popular tool for studying intricacies of molecular processes (Ozsolak and Milos 2011; Conesa et al. 2016; Wang and Ma’ayan 2016; Hasin, Seldin and Lusis 2017). In fact, currently RNA-seq can be considered almost routine if not for the still substantial costs of experiments and subsequent *in-silico* analyses (Svensson, Vento-Tormo and Teichmann 2018), including those associated with data storage and handling (Kwon et al. 2015). This, along with explosive increases in available volumes of data generated in large-scale RNA-seq experiments, contributes to an ongoing demand for universal, easy-to-use computational tools capable of user-specific customization.

One of the widely used workflows available for high-throughput RNA-seq analyses is Galaxy, which is a reproducible and collaborative analytic platform that offers developers a framework for integrating and sharing their tools and workflows (Goecks, Nekrutenko and Taylor 2010; Afgan et al. 2016). Yet, although Galaxy is designed to be relatively easy to use, even for a beginner, performing more in depth analysis with multi-step workflows often requires that a user possesses and/or has access to a specialized bioinformatics expertise. Other challenges are related to sharing potentially large-scale analyses on a public webserver, which can become time-consuming, e.g., with time to completion increasing during high peak usage hours. Further, while there are hundreds of workflows currently accessible on Galaxy, many of these are quite complex and have a substantial learning curve to perform analyses and/or often require user knowledge of reference genomes and file formats. This limits the types of datasets that can be analysed without deploying a custom Galaxy instance, which in turn requires specialized skills. Likewise, for tasks beyond the basic transcriptome discovery analysis the user would need to know how to install and utilize additional tools in the Galaxy instance, somewhat hampering its usability to the potential user with only the basic computing skills. We would like to note that Galaxy Training Network (https://training.galaxyproject.org/) already provides a variety of excellent tutorials to help inexperienced Galaxy users to performed complex analyses (Batut et al. 2018). These tutorials nonetheless require substantial time and effort investments from users, which may exclude small labs lacking necessary manpower or somewhat limit Galaxy’s usability in the classrooms. In the past few years several toolboxes have been released in an effort to address such challenges with using Galaxy (e.g., Grüning et al. 2016; Hung et al. 2016; Meiss et al. 2017; Tithi et al. 2017; Beccuti et al. 2018; Hung et al. 2018). Yet, these toolkits are often designed to analyse only one specific dimension of transcriptome diversity, and/or not fully automated and require some prior knowledge of R command line script (Li et al., 2016).

## Implementation

### AIDD features overview

To help overcome some of these limitations, our pipeline - Automated Isoform Diversity Detector (AIDD) - has been designed implicitly with a novice user in mind, and thus, can be used, for example, as an educational tool for RNA-seq-based laboratory exercises in the classroom setting with a minimal prior user training. Because the pipeline is packaged in a VirtualBox environment, it is easy to install on essentially any operating system and/or a broad range of hardware (Windows, Linux, MacOS) that is capable of handling a VirtualBox installation without concerns for compatibility. Yet despite the seeming simplicity of installing it, our AIDD pipeline is powerful enough to handle a broad range of RNA-seq analyses, spanning from differential gene and isoform expression, to variant calling, and RNA editing analysis using dimension reduction and machine learning approaches, including Guttman scale patterns (Proctor 1970) for time series analysis of ADAR editing landscapes. Unlike comparable tools, AIDD offers a fully automated data analysis pipeline with a simple setup and one-click execution, while still allowing for easily customizable options to account for a wide range of experimental conditions that users may wish to include. AIDD incorporates GATK haplotype caller (DePristo et al. 2011), which is currently not available from Galaxy, as a variant caller for RNA editing prediction, customizable R and bash scripts for detailed statistical analyses of the transcriptome, including RNA editing patterns as well as transcriptome-level differential expression combined with gene enrichment and pathway analysis. SnpEff (Cingolani et al. 2012) is used to add depth to the complete transcriptome analysis by predicting the impact of RNA editing on protein structure and function. AIDD also performs data visualization as part of the automated pipeline and produces publication-ready heatmaps, volcano and violin plots, bar charts and Venn diagrams.

### AIDD availability and hardware requirements

The AIDD pipeline is built in an Oracle VirtualBox (https://www.oracle.com/virtualization/virtualbox/index.html) virtual machine based on Ubuntu 18.04.2 LTS (Bionic Beaver) 64-bit PC (AMD64) desktop image (http://releases.ubuntu.com/18.04/) and contains all tools necessary for transcriptome-level analysis (Figure 1). The distributed VirtualBox image is ∼ 20Gb in size and is publicly available for download via GoogleDrive link (https://drive.google.com/open?id=1XOWh9H-v1nA6_Vl53PI6G2gKaVoZX6ls). The up-to-date detailed description of included software tools, AIDD manual and step-by-step tutorial for AIDD are distributed via our GitHub site (https://github.com/RNAdetective/AIDD).

**Figure 1:**
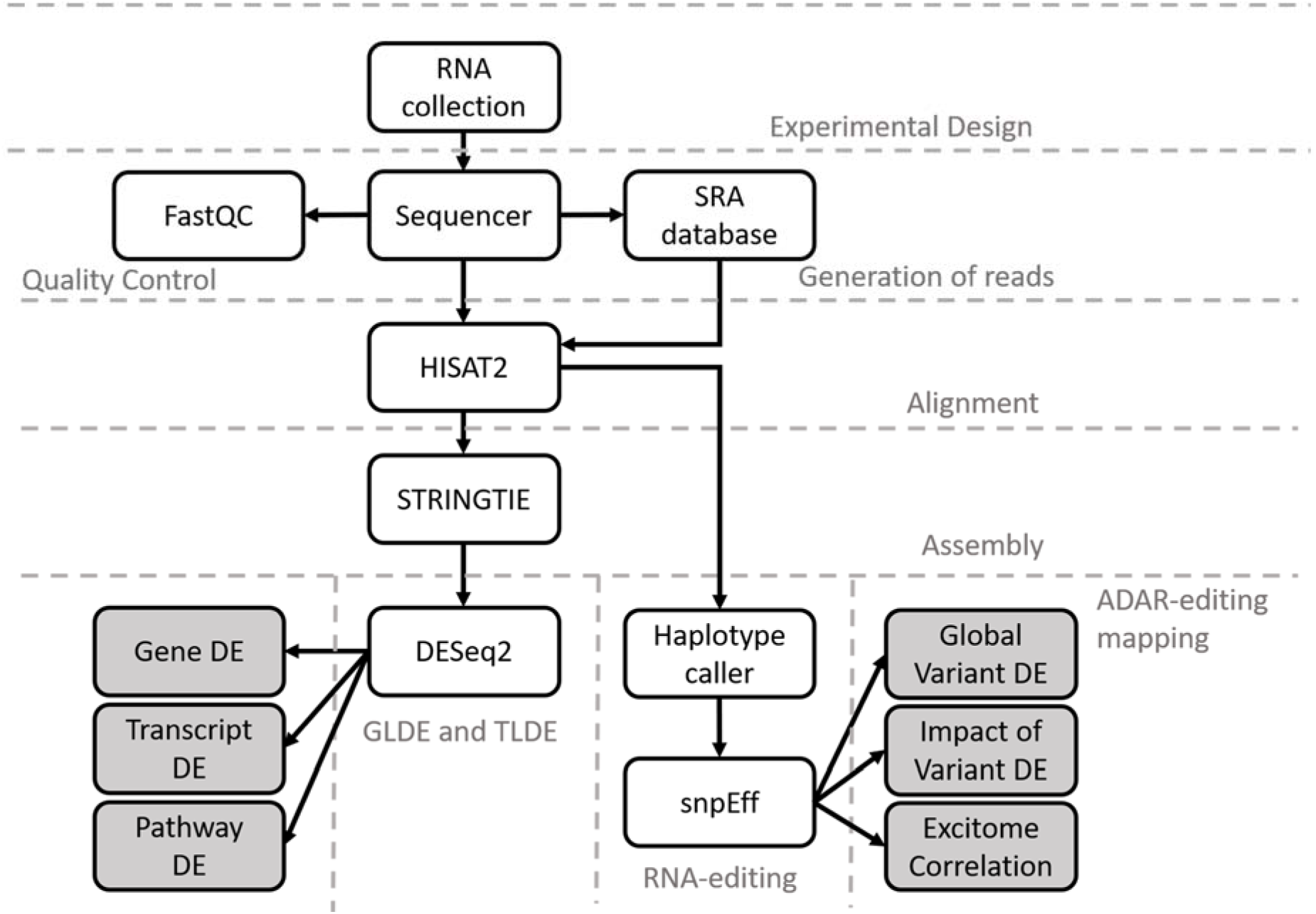
Flow chart of the tools and steps used in the automated workflow carried out by AIDD pipeline. The analysis begins from gathering relevant RNA-seq data files from the NCBI SRA database, followed by reads alignment using HISAT2 with Ensembl annotations. Transcriptome assembly is then performed by Stringtie. Downstream expression analysis can be performed using multiple tools, including DESeq2, edgeR and topGO. Variant calling to detect RNA-editing events, including A-to-I editing, is performed using tools implemented in GATK; and statistical analysis of the effect of RNA editing is performed using custom R scripts.

Implicitly tailored toward a novice user with no or minimal experience in computational analyses, AIDD is designed to run automatically with limited user input through a customizable bash script that controls multiple computational tools, including HISAT2 and GATK, among others, to comprehensively analyse RNA-seq datasets. AIDD can be deployed on almost any modern laboratory, classroom or office computer capable of running Ubuntu 18.04 in a VirtualBox environment. To shortcut the early learning curve, the pipeline is set up to run with default parameters directly “out of the box”, and includes commented out examples in the form of R markdown file that the user can choose to deploy as a step-by-step tutorial.

The minimum recommended hardware specifications include 4 GHz dual-core processor (or better), 8 to 12 GB system memory available to the virtual environment, and 50 GB of free hard drive space (https://www.ubuntu.com/download/desktop), although at least 16 GB system memory is recommended, and some applications may require more. For example, STAR alignment tool needs at least 10 times more memory bytes than the target genome, which for human genome translates into at least 32 GB and upwards if annotations are needed (Dobin and Gingeras 2015).

### Included example datasets: transcriptomes of ZIKV-infected neural progenitor cell lines and importance of ADAR gene family

To illustrate the AIDD capabilities, we use a publicly available dataset from a study by McGrath et al. (2017) that contains RNA-seq data from three genetically distinct neural progenitor cell (NPC) lines infected with Zika virus (ZIKV) (McGrath et al. 2017). The authors found varying degrees of severity of symptoms associated with congenital Zika syndrome (CZS), including decreased differentiation and proliferation, and increased signs of apoptosis (McGrath et al. 2017).

McGrath et al. also reported increased expression of genes involved in innate immune response, including interferon alpha (IFNA) and adenosine deaminase acting on RNA (ADAR) during ZIKV infection (Supplementary Table 1 in McGrath et al. 2017). The ADAR gene family consists of three genes, namely, ADAR (also referred to as ADAR1), ADARB1 (ADAR2), and ADARB2 (ADAR3). Only ADAR and ADARB1 have proven deaminase activity (Chen et al. 2000; Jin, Zhang and Li 2009; Walkley, Liddicoat and Hartner 2011) catalyzing the deamination of adenosine (A) to inosine (I) transition seen in RNA editing (Nishikura 2010; Savva, Rieder and Reenan 2012). ADARB2 is thought to play a regulatory role through competition with other ADARs for substrate binding (Hardt et al. 2008; Savva, Rieder and Reenan 2012). ADARs play a prominent role in the nervous system (Maas, Rich and Nishikura 2003; Tan et al. 2009; Savva, Rieder and Reenan 2012), specifically in the brain (Mehler and Mattick 2007; Liscovitch et al. 2014), where the majority of ADAR editing target genes are expressed (Melcher et al. 1996; Chen et al. 2000; Gonzalez et al. 2011; Li and Church 2013), including during development (Wahlstedt et al. 2009).

### Running AIDD: Uploading RNA-seq data into AIDD

AIDD is designed to automatically download and convert RNA-seq datasets from the SRA accession numbers that user defines in the experimental conditions table. For the example analysis discussed here, a subset of Bioproject PRJNA360845 (McGrath et al. 2017) was downloaded and converted to fastq format. Once converted to fastq format, fastqc (http://www.bioinformatics.babraham.ac.uk/projects/fastqc/) is used for quality control. Upon user assessment of quality of files, fastx-Toolkit (http://hannonlab.cshl.edu/fastx_toolkit/) is used to trim fastq files to assure best quality for alignment. In addition to downloading and preparing sequences, AIDD also automatically downloads and formats all necessary default references and indexes for human genome to run the tools. There are also options for user-defined reference sets, e.g., if RNA-seq data comes from mouse rather than human. AIDD can also run from locally stored fastq or standard alignment SAM/BAM files.

In addition to PRJNA360845 RNA-seq data (McGrath et al. 2017), the included tutorial uses a second dataset from Bioproject PRJNA313294 (Tang et al. 2016). While PRJNA313294-based results are not discussed here, they are available through the AIDD manual and in the distributed AIDD image (https://github.com/RNAdetective/AIDD).

### Running AIDD: Reads alignment and assembly

Once the RNA-seq data and the reference files have been downloaded, the reads are aligned to the chosen reference (GRCh37_snp_tran is used as a default, and in this example). The pipeline uses HISAT2 (Kim, Langmead and Salzberg 2015) as a default alignment tool. SALMON (Patro et al. 2017) and STAR (Dobin et al. 2013) aligners are also available as options. The HISAT2 (https://ccb.jhu.edu/software/hisat2/index.shtml) aligner is a low-memory yet sensitive alignment program that allows for comparable results to other slow and more memory intensive aligners such as STAR (Dobin and Gingeras 2015; Kim, Langmead and Salzberg 2015). Once the reads have been aligned, the output files (SAM format) are converted into BAM format using Picard tools (http://broadinstitute.github.io/picard/) in preparation for variant calling and transcriptome analysis. The pipeline saves these intermediate files should the user ever need to use them for additional analyses.

Next, the transcriptome is reconstructed using Stringtie (Pertea et al. 2015), with cufflinks available as an option (https://software.broadinstitute.org/cancer/software/genepattern/modules/docs/Cuffdiff/7), with output generated as raw counts (Fragments Per Kilobase Million (FKPM), Transcripts Per Kilobase Million (TPM), and coverage) in the “counts” folder, and gene transfer format (GTF) files. The latter are then automatically modified into the count matrix for subsequent input into DESeq2 (Love, Huber and Anders 2014; Varet et al. 2016), using the coverage correction for raw counts unique to Stringtie. The conversion step is performed by a Python script available from the Stringtie website (https://ccb.jhu.edu/software/stringtie/).

### Running AIDD: Differential Expression Analysis

Once reads have been mapped, DESeq2 (Love et al., 2014) and other dependent packages are used to generate gene-level and transcript-level differential expression outputs, including results of the principal component analysis. The latter can be used as a quality control or as an exploratory analysis step, to verify the similarity among samples or treatments, and to identify outliers. DESeq2 uses empirical Bayes shrinkage approach to take into account within-group variation as well as fold change estimation to control for variance observed in the low read count genes (Love et al., 2014). This approach allows for increased sensitivity and decreased false positive rate (Love et al., 2014). A user supplied gene list, for example, a Gene Ontology (GO)-based list, can be used to create pathway expression heatmaps and volcano plots to visualize significantly differentially expressed genes involved in those user-defined pathways, along with the default pathways for GO terms involved in neural development, proliferation, differentiation and signalling as well as the gene list of the innate interferon pathway that we used to explore the role of ADAR editing in CZS (Supplementary Tables 1-5). Additional pathway enrichment analysis is automatically performed using included R package topGO (Alexa and Rahnenfuhrer 2010). Alternatively, generated gene and transcript lists can be used with outside gene enrichment analysis tools such as PANTHER (Mi et al. 2010) or DAVID (Huang da et al. 2007).

### Running AIDD: Variant Calling

While the state of the art identification of genomic variants that can be linked to phenotypic variation is based upon whole-genome (WGS) or whole-exome sequencing (WES) (Piskol, Ramaswami and Li 2013), much broader availability (and affordability) of transcriptome sequencing data makes it another appealing source of variants discovery (Han et al. 2015). Furthermore, some mechanisms of variants generation – such as RNA editing and splice-site variation – can only be studied at the transcriptome level. Thus, our pipeline includes tools enabling variant discovery from transcriptome data, with the focus on ADAR-mediated RNA editing.

GATK haplotype caller (McKenna et al. 2010) is the tool used in AIDD to infer potential RNA editing events, based upon the best practice settings as defined by the GATK developers as of March 2019 (https://software.broadinstitute.org/gatk/documentation/article.php?id=3891). Picard tools are used for quality control and proper formatting of input files. Haplotype caller is used twice in the pipeline, along with filtering steps to control for both false positives and false negatives. SnpEff is then used to predict consequences on protein structure and function for the inferred variants (Cingolani et al. 2012). Once a final list of potential variants is generated, these are then processed using R scripts to demonstrate both global and local view of RNA editing. Additional set of R scripts will then compare differential ADAR editing landscapes between conditions. It should be noted that here we focus on potential editing events within coding regions, and thus, we are not considering hyperediting events (Porath, Carmi and Levanon 2014). Likewise, genomic polymorphisms can appear as potential editing events in RNA-seq, and thus we include an annotation of detected edited site candidates with available polymorphism data (where applicable). Figure 2 and Supplementary Table 6 outline various tools, used, as well as folders and files generated by the pipeline.

**Figure 2:**
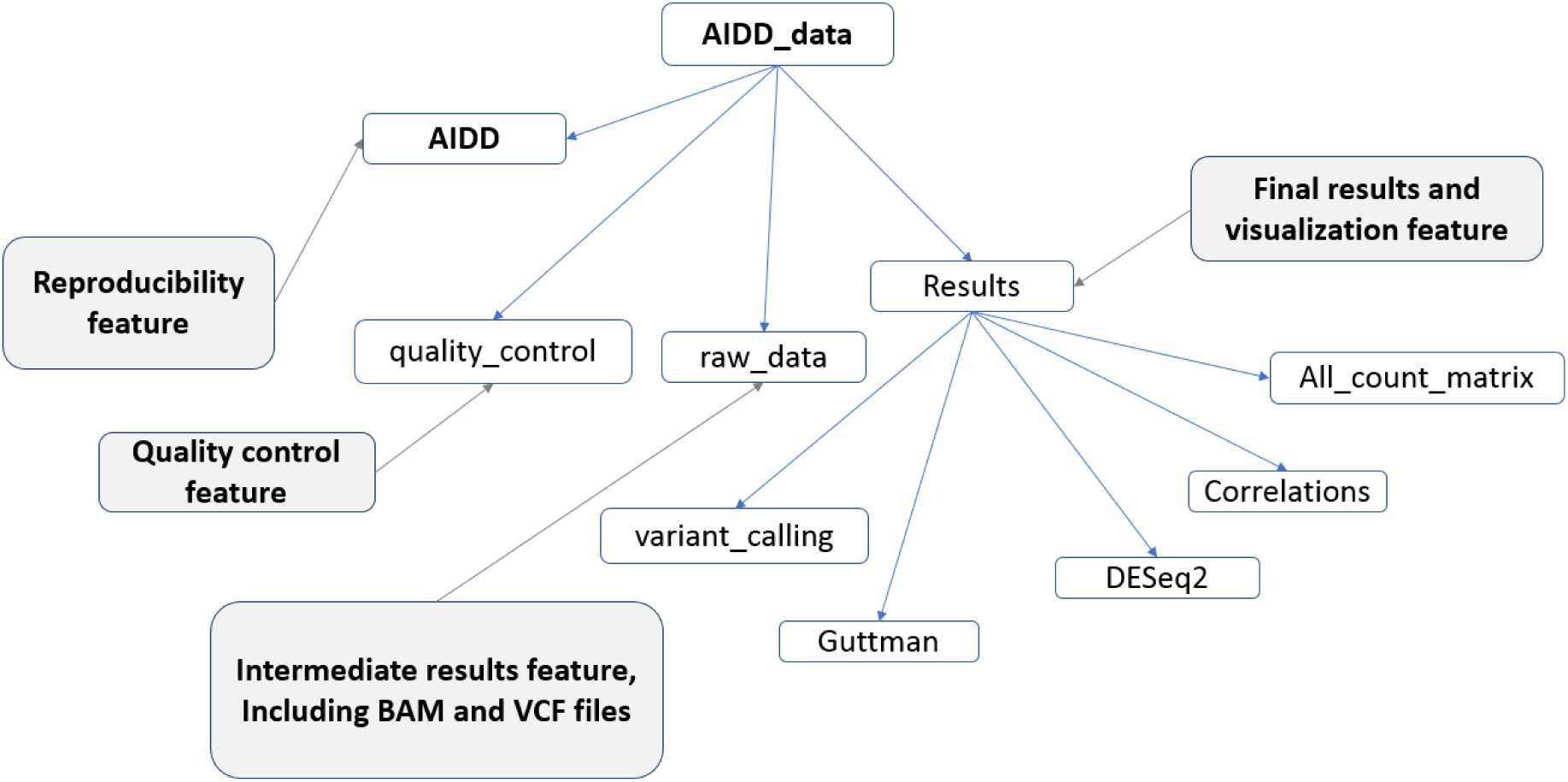
Flow chart showing directory structure created by AIDD. The main folder is AIDD_data and contains 4 folders including (i) AIDD, containing all scripts used in analysis for reproducibility, (ii) quality control files, (iii) intermediate files, including BAM, GTF and VCF files, (iv) results of statistical analysis and data visualization including differential isoform expression and ADAR editing landscapes.

## Results and discussion

To illustrate AIDD’s capabilities, we describe results from the included tutorial that uses Bioproject PRJNA313294 data from (McGrath et al. 2017). Using PRJNA313294 data, AIDD mapped reads and then computed normalized and transformed gene and transcript count matrices for differential expression (DE) analysis using DESeq2 with a multivariate model for infection status taking into account cell-line identity. Principle component analysis (PCA) of the top 500 expressed genes showed that ∼47% of the variance is explained by the first principle component, which separated cell lines by fetal age, with K048 cell line derived from the 9 week old fetal tissue being separated from the 13 weeks old fetal tissue of G010 and K054 cell lines. The second principle component explained ∼27% of the variation, and clustered ZIKV-infected cells from the mock infected cells, except in the case of the G010 cell line (Figure 3A). The pipeline also generated a heatmap of the top 60 differentially expressed genes with hierarchal clustering that showed clustering of samples by infection status, except for the G010 cell line (Figure 3B). This latter phenomenon is consistent with reported findings of McGrath et al. (McGrath et al. 2017) that showed that G010 cells exhibited the least amount of cytopathic effects, if any, due to ZIKV infection, potentially reflecting genetic heterogeneity across studied cells. Figure 3C shows generated volcano plots that visualize the top 20 differentially expressed genes between ZIKV and mock infections taking into account differences in cell-lines. AIDD generates clustering heatmaps for each cell line, which showed that while both K048 and K054 exhibit clear differences between mock and ZIKV infections consistent with the phenotypic differences between the two conditions (Figure 3D & E), G010 cells showed no significant difference between ZIKV and mock infected cells, consistent with McGrath et al. (2017) results (Figure 3F). By looking at each cell line individually, AIDD is able to highlight differential effects of ZIKV infection in combination with host genetics that are consistent with results originally reported by McGrath et al. (2017) (Figure 3G, H & I).

**Figure 3:**
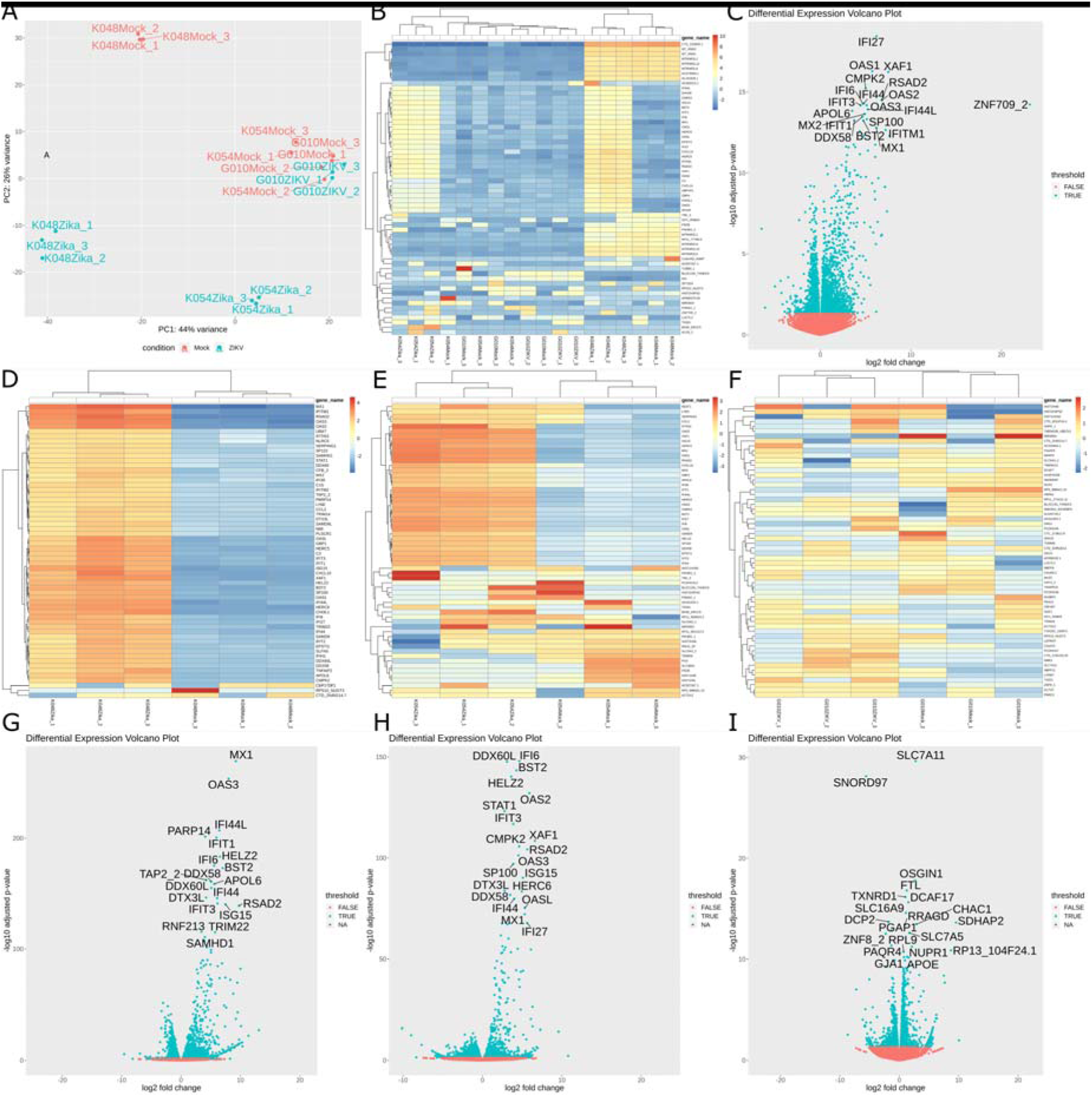
Visualization of differential expression analysis using AIDD. (A) Principle component analysis of the top 500 expressed genes counts show 47% of the variance in the system is attributed to differences in cell lines and 27% of the variance is attributed to ZIKV infection status. (B) The top 500 hierarchal clustering also shows clustering of CSZ phenotype cell line (K048 & K054) ZIKV infected cells and normal phenotype cells (G010) regardless of ZIKV infection status clustered with the CSZ phenotype cell line mock infections. (C) The top 20 differentially expressed genes during ZIKV infection taking into account genetic cell line differences highlight the innate immune activation. When looking at each cell line independently, K048 (D) and K054 cells (E) have clear pattern of differentially expressed genes during ZIKV infection, whereas G010 cells (F) shows less of a pattern of differentially expressed genes. Panels G-I show that when the top 20 differentially expressed genes are considered, each genetically distinct cell line shows a differentially gene expression response to ZIKV infection.

### Pathways analysis

The gene pathways exploration tool included in AIDD was used to examine differential expression in neurodevelopmental pathways during ZIKV infection. Using gene list supplied by the user, AIDD will generate customized heatmap, volcano plot, and data table with differential expression results for genes of interest. Gene ontology (GO) terms “innate immunity”,” brain development”, “central nervous system development”, “neurological development”, and “peripheral nervous system” are already included as default pathways. We also included a custom gene list for genes in the interferon alpha pathway (Supplementary Table 1). AIDD results showed that ZIKV-infected cells showed increased expression of innate immune genes (Figure 4A), as well as those in the interferon alpha pathway, including ADAR (Figure 4B), except for the G010 cells. Consistent with McGrath et al. findings (McGrath et al. 2017), cell lines that have CZS-like phenotypic appearance if ZIKV infected (namely, K048 and K054) have significant differential expression in the majority of the genes involved in the interferon alpha pathway (Figure 4C & D), whereas G010 cells that appear to be essentially normal phenotypically showed only a few significantly differentially expressed genes in the interferon alpha pathway (Figure 4E), pointing to potential involvement of interferon alpha response in ZIKV infection and CZS-like symptoms (Piontkivska et al. 2019). On the other hand, only cell line-associated differences but not the ZIKV infection-mediated differences were observed for genes associated with GO terms of brain development (Figure 4F), central nervous system development (Figure 4G), neurological development (Figure 4H), and peripheral nervous system development (Figure 4I).

**Figure 4:**
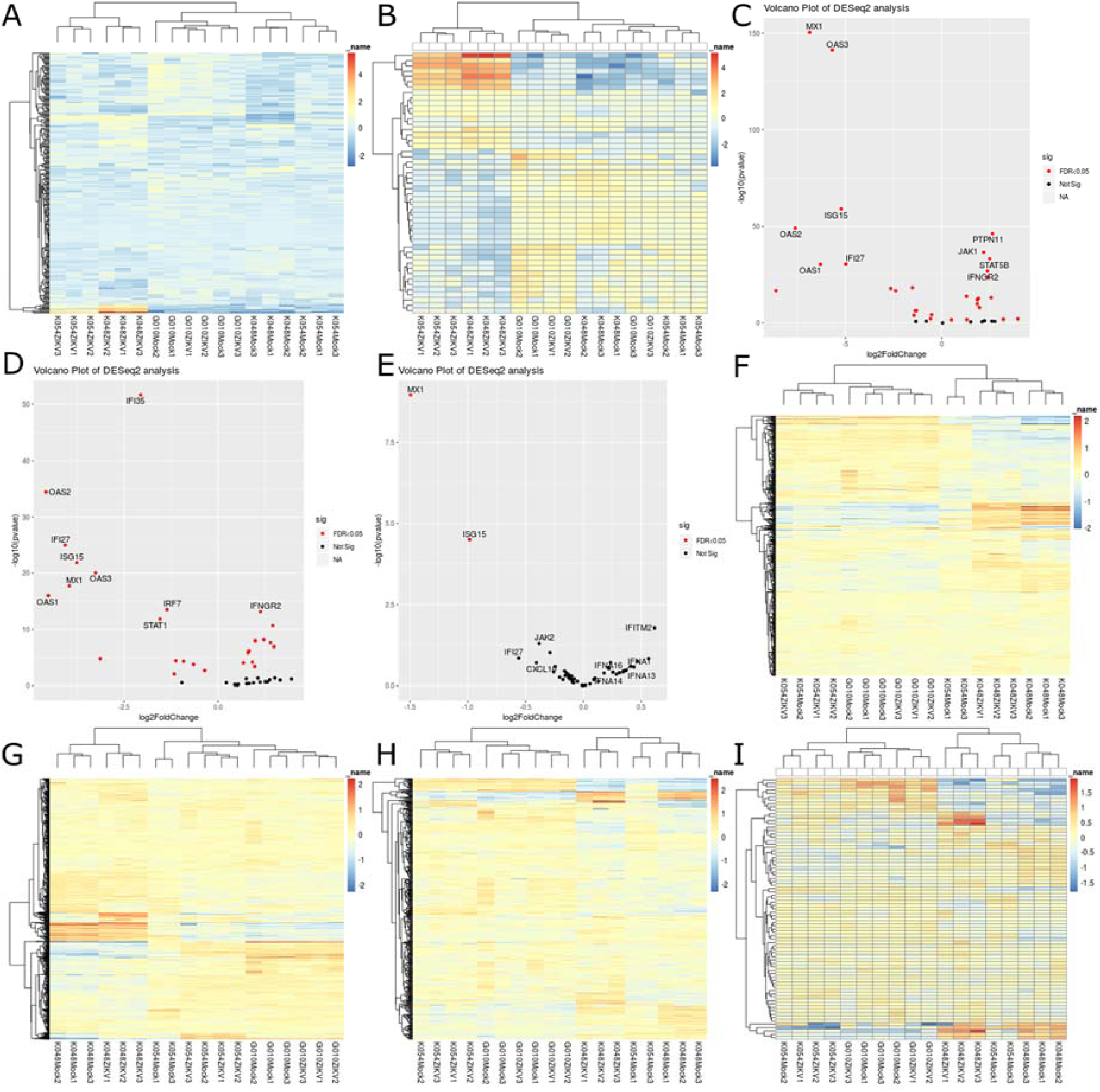
Results of AIDD pathway expression analysis. (A) Gene Ontology term “innate immune system” shows clustering of ZIKV infected cells with the CSZ phenotype (K048 & K054) and clustering of normal phenotype (G010) with the mock infected cells of all 3 cell lines. (B) Customized “interferon alpha pathway” list shows similar clustering pattern as (A). The CZS phenotype cell lines (K048 & K054) show the top 10 differentially expressed genes with gene products induced by interferon alpha pathway, including OAS1 and 2, and intermediary genes in the interferon alpha pathway, including STAT1 (C & D, respectively). On the other hand, phenotypically normal cell line (G010) has only 2 differentially expressed genes, which are not part of the interferon alpha pathway (E). Gene ontology terms “brain development” (F), “CNS development” (G), “neurological development” (H), and “PNS development” (I) exhibit differential expression patterns that can be attributed to genetic differences among cell lines, but not associated with ZIKV infection.

### Mapping ADAR expression and editing landscapes

To explore the potential role of ADAR enzymes and ADAR editing, AIDD allows us to focus on expression of ADAR genes and editing patterns (Supplementary Tables 7 & 8), including applying Guttman scale patterns to identify temporal changes in ADAR editing landscapes (Supplementary Figure 1). The results showed that ADAR1p150 isoform-specific expression was significantly higher in ZIKV infected cells with the CZS phenotype (K048 and K054), while not being significantly different in G010 cells (Figure 5A). Interestingly, ADARB1 showed the opposite pattern, being significantly upregulated in G010 cells, but not in cells with CZS-like phenotype (Figure 5B). Because ADARB1 and ADAR both share some overlapping editing targets as well as have gene-specific ones (Lehmann and Bass 2000; Riedmann et al. 2008), this expression pattern suggests that both ADAR genes may play complementary roles in the differential response to ZIKV infection (Piontkivska et al. 2019). This would be consistent with prior suggestions that ADARB1 contributes to dysregulation of RNA editing in many diseases (Amore et al. 2004; Cenci et al. 2008; Hideyama et al. 2012; Karanovic et al. 2015).

**Figure 5:**
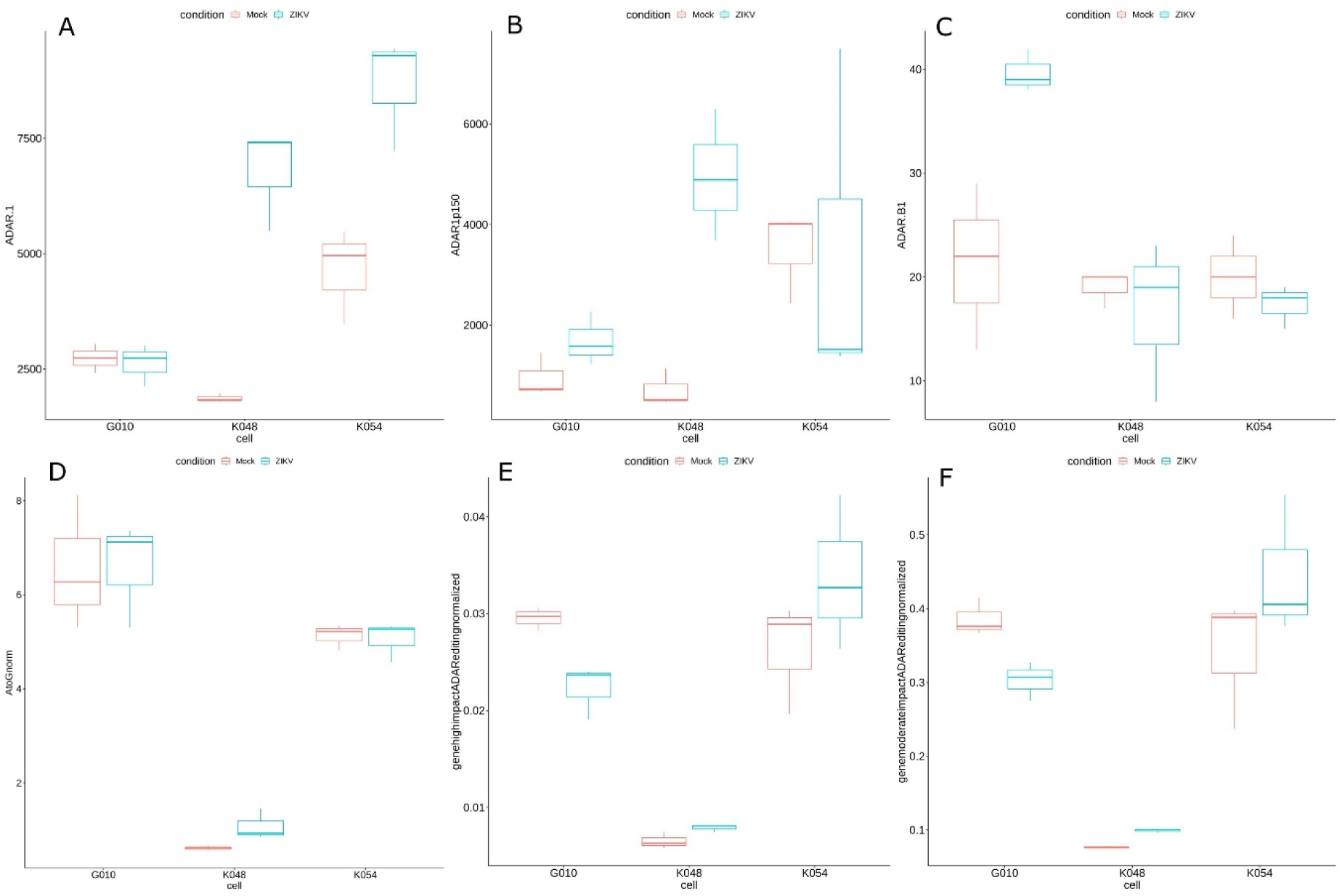
Visualization of ADAR expression and ADAR editing landscapes. (A) ADAR expression is significantly increased in CZS phenotype cell lines K048 (F=58.396, p=0.001575) and K054 (F=18.516, p=0.01261), but not in phenotypically normal G010 (F=0.1219, p=0.7446) cells. (B) ADAR1p150 expression is significantly higher in K048 (F=29.497, p=0.005576), but not in K054 (F=2e-04, p=0.9902) or G010 (F=3.4772, p=0.1357) cells. (C) ADARB1 expression is not significantly different in K048 (F=0.2579, p=0.6383) or K054 (F=1.0492, p=0.3636) cells, but is significantly higher in G010 (F=14.684, p=0.01859). (D) The numbers of A to G substitutions were somewhat elevated in K048 (F=6.0422, p=0.06984), but not in G010 (F=6e-04, p=0.9813) or K054 cells (F=0.0648, p=0.8116). (E) The numbers of A to G substitutions with predicted high impact on protein structure and function were significantly lower in G010 (F=17.498, p=0.01388), but somewhat higher in K048 cells (F=6.3489, p=0.06538); there were no changes in K054 cells (F=1.7384, p=0.2578). Likewise, moderate impact substitutions were also significantly lower in G010 (F=15.737, p=0.01658) and significantly higher in K048 (F=157.23, p=0.0002328) cells, while were not changed in K054 cells (F=1.9198, p=0.2381) (F).

AIDD also allows the user to map ADAR editing landscapes by performing variant calling to identify potential A to G substitutions. Globally, we found that the total numbers of A to G substitutions are higher in ZIKV-infected in both the G010 and K048 cell lines but not in the K054 line (Figure 5C). However, when the potential impact of these substitutions on protein structure and function is examined, cell lines with the CZS-like phenotype (K048 and K054) had more of high and moderate impact variants detected in ZIKV infection, while seemingly normal G010 cells had smaller number of potentially impactful changes in ZIKV infection (Figure 5D, E & F).

It should be noted that one major challenge of using variant calling methods for detecting RNA editing events is the need to have a sufficient coverage depth (of at least 50 million reads or higher per sample) to accurately detect editing events when editing frequencies are low. AIDD attempts to correct for this by normalizing substitution counts by the read depth as determined from alignment algorithms. Therefore, these observed editing differences among cell lines could be attributed to interactions between ADAR family members as well as ADAR preferences at the editing sites, and spatio-temporal regulation of editing.

We were also interested in editing events at known editing sites in ion channels and transporters that are known to be associated with fine-tuning of neural signalling, including excitotoxicity, brain development and neural plasticity (Tan et al. 2009; Hood and Emeson 2012; Eran et al. 2013). To define the excitome, computationally-predicted ADAR editing sites found in psychiatric disorders confirmed with PCR (Zhu et al., 2012) were combined with editing sites from RADAR database that were previously examined in Alzheimer’s disease (Khermesh et al., 2016) to create a list of 151 editing sites located in 91 genes (Supplementary Table 8). In part because of relatively low coverage in all three cell lines as well as rather drastic differences in fetal age, the editing patterns at specific sites varied both between different cell lines and between infected and uninfected cells. ZIKV infected K048 cells showed likely editing events at multiple sites, including at two ion channel receptors (namely, GRIA3 and GRIN3B). Other ZIKV-induced editing events were detected at IGFBP7, KIF20B and SRP9 genes, responsible for controlling cellular metabolism, vesicular transport, and proper protein storage and transport respectively (Godfried Sie et al. 2012; Ivanova et al. 2015; Lee et al. 2017; McNeely, Little and Dwyer 2019). There was also an increased editing detected at the ATXN7 gene that is implicated in degenerative ataxia (Clark et al. 2015). ZIKV infected K054 cells showed likely editing events in PTPRN2, GRIA2 Q/R site, GRIA3 and IGFBP7, whereas uninfected cells showed editing events in ATXN7, BEST1, BLCAP, and KIF20B. ZIKV infected G010 cells exhibited increased editing in ATXN7, KIF20B, and PTPRN2, and decreased editing at the NEIL1 genes. Changes in editing landscapes can also be visualized with Guttman scale patterns, where differences between distinct cell lines as well as mock and infected cells are shown for individual editing events/residues (Supplementary Figure 1). However, further transcriptomics studies – including at much higher read depth - are needed to fully elucidate the changes in editing patterns that can be induced by viral infections.

## Conclusions

A fully automated pipeline, Automated Isoform Diversity Detector (AIDD), has been developed to facilitate RNA-seq analyses focused on changes in transcriptome diversity, including isoform expression ratios and ADAR-editing events. A publicly available dataset of human neural progenitor cells (McGrath et al. 2017) is used to demonstrate how AIDD pipeline can be used to robustly and reproducibly analyse transcriptome diversity and to infer RNA editing patterns from RNA-seq data. Presented results illustrate the importance of examining both the gene-level and the isoform-level expression differences, as well as exploring RNA editing aspects of transcriptome diversity and their potential association with pathogenicity mechanisms.

AIDD pipeline has additional benefits of being novice-user friendly and completely automated for highly reproducible results. Briefly, AIDD incorporates multiple steps needed for using RNA-seq data to study transcriptome diversity, and offers an easy-to-use pipeline for mapping and contrasting genome-wide RNA editing patterns, with focus on protein-coding transcripts (Supplementary Table 8). Once reads have been mapped to the reference genome, AIDD uses DESeq2 to infer patterns of differential expression at both gene and transcript levels. For users - such as ourselves - interested in patterns of editing of excitome-related genes, AIDD will summarize the expression of the excitome gene members, including ADARs and other genes with known editing sites. AIDD will further summarize global RNA editing patterns and infer correlations between edited sites and ADAR expression patterns. Lastly, lists of genes involved in ADAR editing landscape changes are produced and can be used as potential biomarkers for diagnostic and prognostic purposes.

The distributed pipeline image includes a user-friendly tutorial written in R markdown that can be used to illustrate AIDD features in a classroom setting as teaching tool and/or to generate hypotheses for future experimental validation, or both. The ZIKV infection-associated example described in this paper further highlights the ability of AIDD to conduct complicated analyses within the constraints of a small research laboratory. Future work includes testing AIDD’s accuracy against simulated reads with known editing sites and across various read depths per sample, as well as expanding AIDD’s ability for variant calling by incorporating other methods (such as Freebayes, Garrison and Marth 2012). AIDD can also be used in meta-analysis of publically available RNA-seq datasets to comprehensively map ADAR editing landscapes across different cells and organisms, and to facilitate discovery of novel diagnostic and prognostic biomarkers and potential targets for drug therapies.

## Declarations

### Ethics approval and consent to participate

Not applicable.

### Consent for publication

Not applicable.

### Availability of data and materials

The datasets used in this current study are publicly available in the NCBI SRA/BioProject repository, at https://www.ncbi.nlm.nih.gov/bioproject/PRJNA360845/ and https://www.ncbi.nlm.nih.gov/bioproject/?term=PRJNA313294.

The AIDD pipeline is distributed via GitHub, at https://github.com/RNAdetective/AIDD.

### Authors’ contributions

NMP designed and implemented the pipeline, and wrote the manuscript. EJ, MF, HM and GF contributed to conceptualization of pipeline features, testing of code components and validation, and provided manuscript feedback. RM and GC contributed to conceptualization of pipeline features and analysis steps. HP conceived the pipeline, supervised the project, helped with code and testing, and wrote the manuscript. All authors read and approved the final manuscript.

## Acknowledgements

Not applicable.

---

Supplementary Tables are available at GitHub, https://github.com/RNAdetective/AIDD/tree/master/AIDD_supplFiles.ST1-8_and_SF1

**Supplementary Figure 1:**
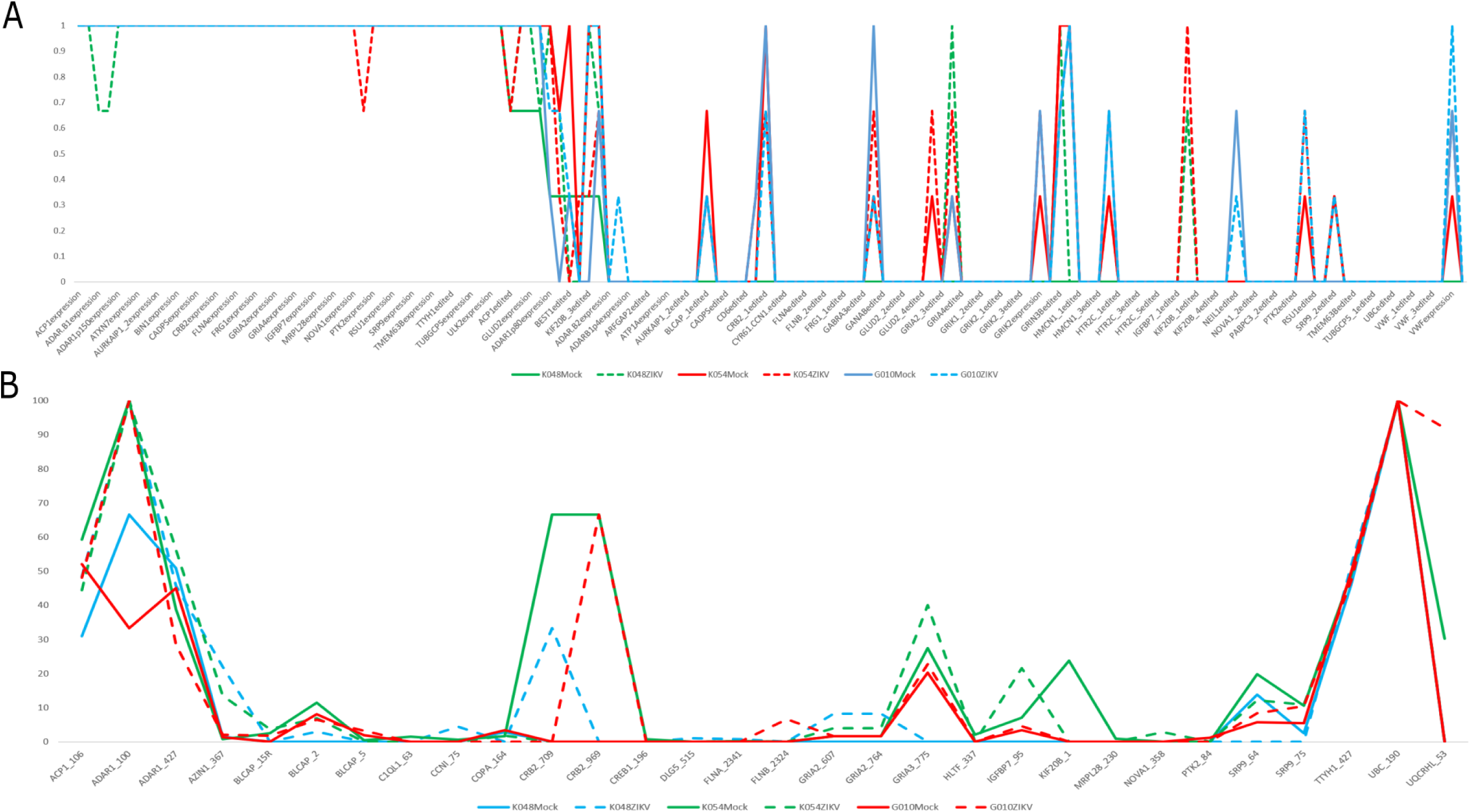
Guttman scale patterns (Proctor 1970) were used to order and group ADAR editing sites based on the frequency of samples that had editing at those sites. ADAR editing landscapes are differentially edited in both order and groupings based on cell line and ZIKV infection. (A) The expression and editing events are ordered by normal phenotype cell line G010 shown in blue, with cell lines K048 and K054 shown in green and red, respectively. The mock-infected cells are shown with solid lines and ZIKV-infected cells are shown with dashed lines. (B) The mean editing frequencies differ between mock- and ZIKV-infected cells at several sites including; (i) AZIN1 at amino acid position 367 (F=7.1095, p=0.00263), (ii) CRB2 at amino acid position 969 (F=3.2, p=0.04584), (iii) IGFBP7 at amino acid position 95 (F=40.651, p=4.09e-07), (iv) SRP9 at amino acid position 75 (F=3.5131, p=0.03459), and (v) UQCRHL at amino acid position 53 (F=8.796, p=0.00105). Changes in editing patterns were also detected at ADAR1 at amino acid position 427 (F=2.9571, p=0.05749), CCN1 at amino acid position 75 (F=2.5546, p=0.08504), and GRIA3 at amino acid position 775 (F=2.5515, p=0.08531), respectively.

