## SupplementaryFiles for "Automated Isoform Diversity Detector (AIDD): A pipeline for investigating transcriptome diversity of RNA-seq data": AIDD.SupplFigure1.pdf

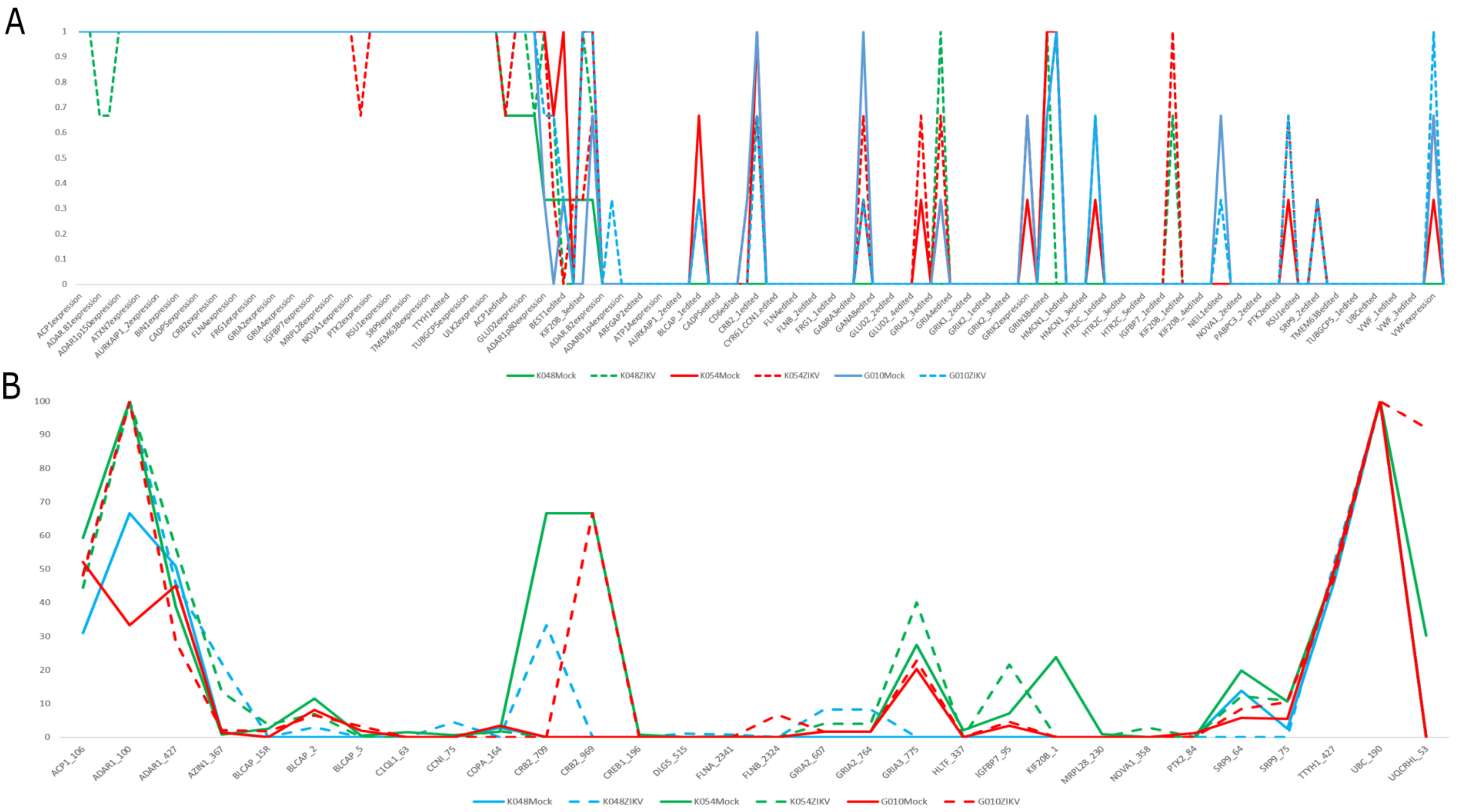


Supplementary Figure 1: Guttman scale patterns (Proctor 1970) were used to order and group ADAR editing sites based on the frequency of samples that had editing at those sites. ADAR editing landscapes are differentially edited in both order and groupings based on cell line and ZIKV infection. (A) The expression and editing events are ordered by normal phenotype cell line G010 shown in blue, with cell lines K048 and K054 shown in green and red, respectively. The mock-infected cells are shown with solid lines and ZIKV-infected cells are shown with dashed lines. (B) The mean editing frequencies differ between mock- and ZIKV-infected cells at several sites including; (i) AZIN1 at amino acid position 367 ($F=7.1095$, $p=0.00263$), (ii) CRB2 at amino acid position 969 ($F=3.2$, $p=0.04584$), (iii) IGFBP7 at amino acid position 95 ($F=40.651$, $p=4.09e-07$), (iv) SRP9 at amino acid position 75 ($F=3.5131$, $p=0.03459$), and (v) UQCRHL at amino acid position 53 ($F=8.796$, $p=0.00105$). Changes in editing patterns were also detected at ADAR1 at amino acid position 427 ($F=2.9571$, $p=0.05749$), CCN1 at amino acid position 75 ($F=2.5546$, $p=0.08504$), and GRIA3 at amino acid position 775 ($F=2.5515$, $p=0.08531$), respectively.
