## SupplementaryFiles for "Automated Isoform Diversity Detector (AIDD): A pipeline for investigating transcriptome diversity of RNA-seq data": AIDD.SupplTable6.docx

**Supplementary table 6. Quality control and other features implemented in the AIDD pipeline (https://github.com/RNAdetective/AIDD).**

| **Feature** | **Tool(s)** | **Description** |
| --- | --- | --- |
| Easy to use | Custom bash, python and R scripts | A set of custom script will produce publication ready graphs from raw fastq files with little or no user input. |
| Reproducibility | Custom bash, python and R scripts | All and any scripts and options used for the analysis are copied and stored in a separate folder as part of the AIDD output. |
| Quality control | Fastqc  Stringtie  DESeq2-rlog and vlog  Principle component analysis (PCA) as part of differential expression inference (DESeq2) | Determines trimming start and end points for the reads.  Normalizes count matrix by taking into account read depth at each site.  Transforms count matrices using default transformation rlog.  Checks for batch effects in the samples and summarizes the variation between samples. |
| Intermediate Files | HISAT2  Stringtie  GATK (haplotypecaller)  snpEff | HISAT2 with Stringtie is used to count gene and transcript expression.  Variant calling is used in combination with snpEff to find editing sites and predict their effect on protein structure and function.  Normalized count matrix by taking into account read depth at each site. |
| Statistical Analysis | Custom ExToolset | R scripts with bash wrappers are used for statistical analysis. ANOVA is used for differential expression analysis. Pearson’s correlation is used to determine which ADAR is responsible for editing and DESeq2 is used for differential expression analysis. |
| Data Visualization | Custom ExToolset | R scripts with bash wrappers used for statistical analysis. Boxplots, bar graphs and scatterplots highlight ADAR editing landscapes. Venn Diagrams, heatmaps, and volcano plots show results of differential expression analysis. |
